# Anatomical identification of the neuroendocrine system in the *Nothobranchius furzeri* brain

**DOI:** 10.1101/2020.10.15.342014

**Authors:** Eunjeong Do, Yumi Kim

**Author notes:** **Grant information:** Institute for Basic Science (IBS-R013-D1).

## Abstract

The hypophysis functions as a central gland of the neuroendocrine system for regulating fundamental body physiology. Upon aging, several hormones produced by the endocrine system are dramatically altered. Recently, *Nothobranchius furzeri* (the turquoise killifish) has become a popular model for aging studies because of its short lifespan and highly conserved aging phenotypes. However, the anatomical details of the major neuroendocrine system of the killifish have not been investigated so far. In this study, we have identified the pituitary and pineal glands of the turquoise killifish, which are critical components of the brain endocrine system. These two neuroendocrine glands were weakly attached to the main body of the killifish brain. The pineal gland was located on the dorsal part of the brain, while the pituitary gland was located on the ventral part. Brain sections containing pineal and pituitary glands were performed and revealed that cells in both the pituitary and pineal glands are densely placed than any other regions of brain. Further, three-dimensional images both in pineal and pituitary glands were uncovered their distinctive cellular arrangements. Vasopressin intestinal peptide (VIP) was strongly expressed in the neurohypophysis of the pituitary gland. Glial cells were found inside the pineal gland, while astrocytes covered the outside. These findings illustrate basic features of the neuroendocrine system of *Nothobranchius furzeri*.

## Introduction

The endocrine system consists of multiple glands and organs that secrete hormones essential for organismal development, reproduction, or homeostatic regulation. The hypothalamo–hypophysis axis, also called the neuroendocrine system, is a master link between the central nervous system and the endocrine system. The pituitary gland governs the secretion of major hormones that maintain body homeostasis such as thyroid-stimulating hormone (TSH), prolactin (PRL), somatolactin (SL), growth hormone (GH), α-melanocyte-stimulating hormone (MSH), β-endorphin, adrenocorticotropic hormone (ACTH), follicle-stimulating hormone, luteinizing hormone, somatolacotropes (Sl), and somatolactin (SL). Based on Green’s nomenclature (Green, 1951), the pituitary gland consists of two major parts, the adenohypophysis (AH, consisting of Rostral pars distalis (RPD), Proximal pars distalis (PPD), and Pars intermedia (PI)) and neurohypophysis (Pars nervosa (PN)). The pineal gland is another critical endocrine system in the brain. The pineal gland primarily secretes melatonin, which is involved in the control of the circadian rhythm (Wurtman et al., 1963). It is known that some of the pineal and pituitary gland hormones are dysregulated during the aging process (Bartke and Darcy, 2017; Frohman, 1994; Muller-Fielitz et al., 2017; Seraphin et al., 2008; Uenoyama et al., 2018).

*Nothobranchius furzeri* (turquoise killifish) is a teleost fish originating from a temporal pond around Mozambique and Zimbabwe. The turquoise killifish is a useful animal model because it is easily bred in captivity and has a relatively short life span (median life span, 9–26 weeks) (Kim et al., 2016; Terzibasi et al., 2008). Despite their short lifespan, the turquoise killifish has a highly conserved aging physiology relating to the neuroendocrine system, including neurodegeneration, decreased fecundity, and cognitive impairments. While the brain anatomy of the turquoise killifish has been previously reported, the neuroendocrine system has not yet been characterized (D’Angelo, 2013). The anatomy of the endocrine system has been well-described for other teleost fish models, such as zebrafish and medaka (Ralph Anken, 1998; Wullimann et al., 1996). Thus, we identified two major neuroendocrine organs in the turquoise killifish brain, which were compared with the zebrafish and medaka endocrine systems as a model for the endocrinology of aging.

## Results

### The gross brain anatomy of the killifish revealed weakly-attached subregions in the dorsal and ventral parts of the brain

The gross brain anatomy of the turquoise killifish brain has been reported previously (D’Angelo, 2013). The turquoise killifish brain comprises the 1) olfactory bulbs, 2) telencephalon, 3) diencephalon, 4) optic tectum, 5) cerebellum, and 6) rhombencephalon. In addition to this six main parts, we found two additional bulging structures in the ventral and dorsal regions of the turquoise killifish brain (Figure 1). The two bulging organs, in particular the ventral one, were weakly attached to the main body of the brain. Thus, it was easy to lose them during dissection and further processing. One of the bulging organs was located in the dorsal brain nearest to the telencephalon and between the optic tectum, measuring approximately 0.4 mm in diameter. The other bulging organ was located in the ventral brain at the anterior part of the hypothalamus and caudal to the optic nerves, measuring approximately 0.7 mm in diameter. In comparison with the brain anatomy of the zebrafish and medaka, these additional bulging organs appeared to correlate with the pineal and pituitary glands. Similar to the dorsal bulging organ of the killifish, the pineal gland of the zebrafish and medaka is located between the telencephalic area and the optic tectum of the dorsal brain. The pituitary gland of the zebrafish and medaka is attached to the ventral brain near the hypothalamus, similar to the second bulging organ observed in the killifish ventral brain. Therefore, the location of the killifish brain hypophysis is similar to the position of that in the medaka.

**Fig. 1.**
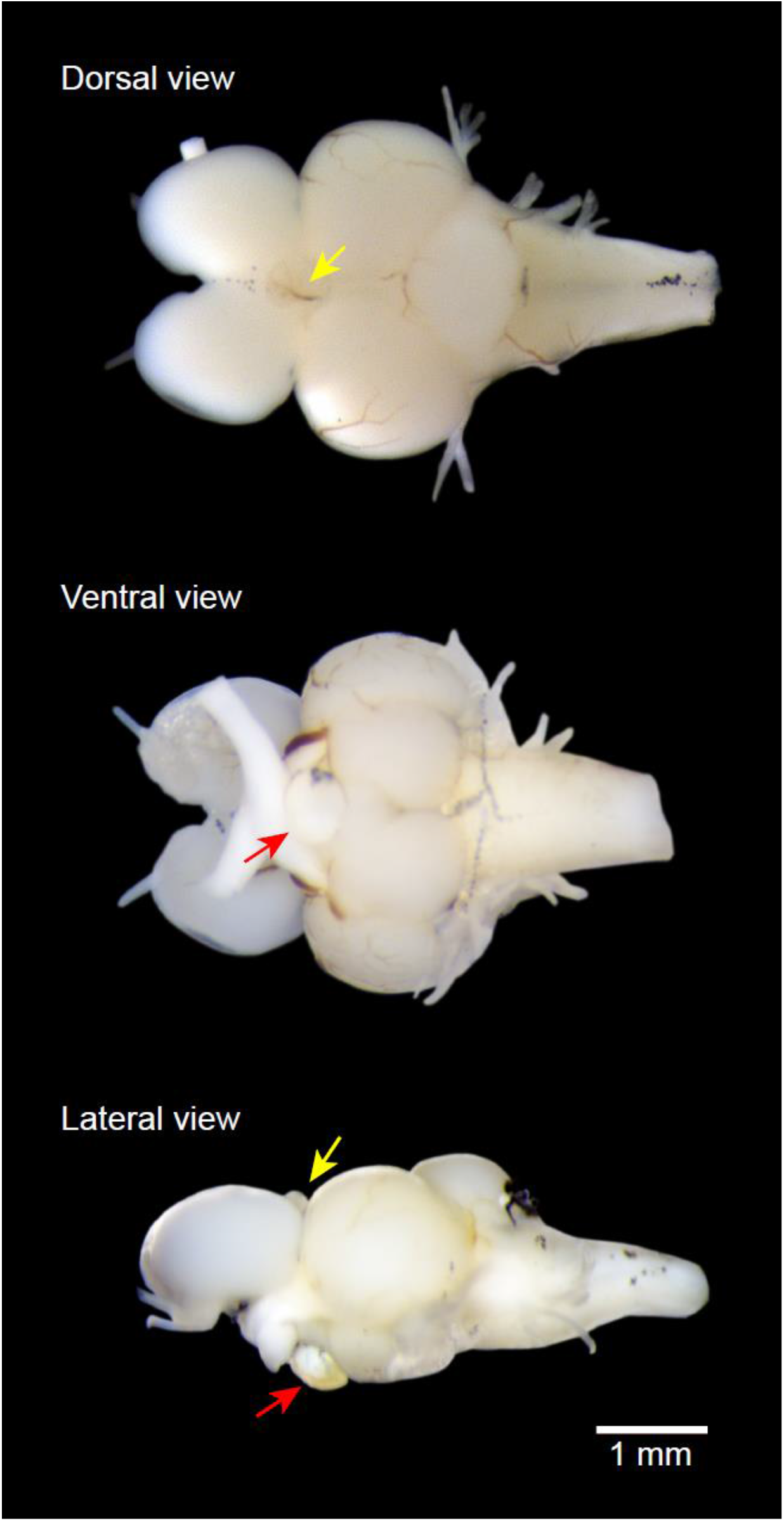
Gross anatomy of the turquoise killifish. Yellow and red arrows indicate pineal and pituitary glands, respectively.

### Characterization of the neuroendocrine organs by Nissl staining

To further characterize the pineal and pituitary glands, a whole killifish brain was sectioned and stained with Cresyl violet (Nissl staining). The internal structures from the brain sections were re-identified based on previous descriptions of *Nothobranchius furzeri* (D’Angelo, 2013), medaka (Ralph Anken, 1998), and zebrafish (Wullimann et al., 1996) brain anatomies. The GRZ-AD, a strain showing the shortest lifespan (16 weeks of median lifespan in our laboratory condition), was used for coronal sectioning of the brain from rostral-to-caudal. With intensive staining, the pineal gland was found at the end of the telencephalon and extended until the beginning of the optic tectum. Sections of the pineal gland was showed a rounded triangular shape at its anterior end (Figure 2a, 2b), then becoming more circular form (Figure 2c), and protruding like a rod from distal end of the pineal gland (Figure 2c). At the location of the distal pineal gland, the pituitary started to emerge and was observed as a separate, flat, and rounded shape. Similar to the pineal gland, the pituitary gland was strongly stained with Cresyl violet compared with other parts of the brain section (Figure 2c). The pituitary bridged the left and right parts of the anterior hypothalamic regions in the diencephalon (Figure 2d-f). The pineal and pituitary glands of the killifish brain could be viewed in one section depending on the sectioning angle (Figure S1). The other parts of the killifish brain, including olfactory bulbs, telencephalon, the remaining optic tectum, cerebellum, and rhombencephalon, are annotated in an illustration (Figure S2, S3).

**Fig. 2.**
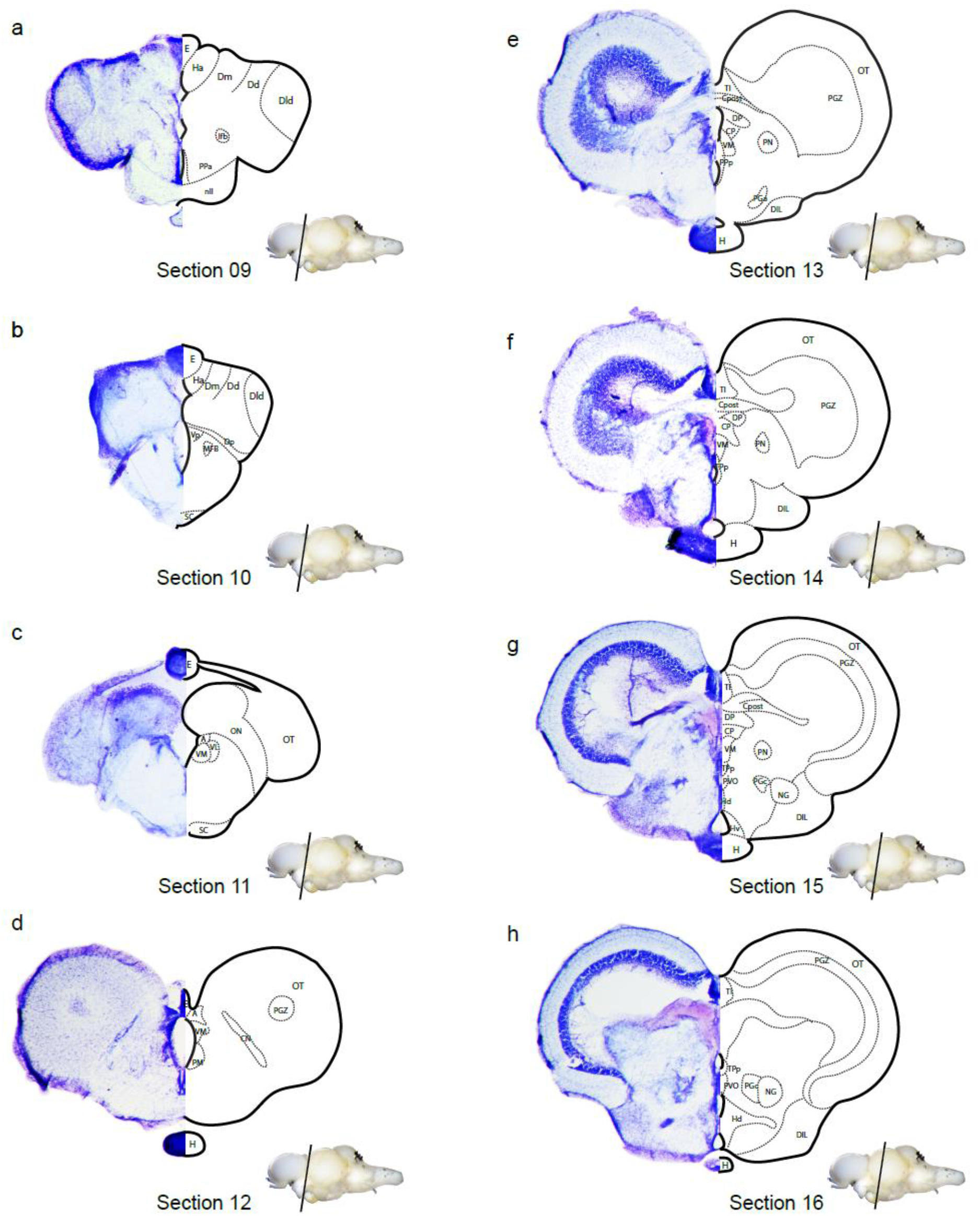
Cross-section of the brain containing the pineal and pituitary glands. **a-d** Sections depict the epiphysis on the dorsal side of the brain. **d-h** Sections depict the hypophysis on the ventral side of the brain. Section numbers are continuous with Supplemental figures 2 and 3. Abbreviations of each figure can be found in the Glossary of brain sections in the materials and methods.

### Anatomy of the turquoise killifish pituitary gland

A 3D imaging method was used to further analyze the detailed structures of the two neuroendocrine glands in the killifish brain. The whole brain tissue was made being transparent and stained with DAPI to observe the intact cell arrangement and internal structure of these glands. As a master gland of the endocrine system, the pituitary gland resembled a flattened sphere covered with blood vessels at the ventral part of the brain (Figure 3a, Supplemental movie 1). To compare the internal anatomy of the killifish pituitary with other teleost fishes’, cell densities were compared from mid-sagittal optical sections. The turquoise killifish exhibited an asymmetric cell density in the pituitary structure, whereby the anterior was more compact than the posterior pituitary gland. The anterior pituitary of the killifish correlated with the RPD, whereas the posterior hypophysis correlated with the PPD, PN, and PI (Figure 3b). The RPD occupied approximately one third of the volume of the killifish pituitary, and cells in this area were arranged compactly. The PPD was located in the ventral pituitary, near to the RPD, PN, and PI. The PN was located in the dorsal pituitary and was surrounded by the RPD, PPD, and PI. Cells in the PN formed clusters surrounded by blood vessels. Additionally, the PN was connected to the hypothalamus as a “Y-shaped” of the turquoise killifish brain (Figure 3c), and a small rounded cavity was observed between the PN and hypothalamus. The PI was positioned in the ventral pituitary. The surface of the pituitary was covered by blood vessels, especially the posterior part of the pituitary gland (Figure 3a). We conducted further analysis by staining with a Vasopressin intestinal peptide (VIP) antibody due to its role in regulating the release of PRL and growth hormone (Kulick et al., 2005; Sherwood et al., 2000). Interestingly, the prolactin-releasing cells are known to be located in the RPD of the medaka (Aoki and Umeura, 1970). VIP staining was mostly detected in the PN. Blood vessels partly covered the RPD of the hypophysis (Figure 3d, Supplemental movie 2).

**Fig. 3.**
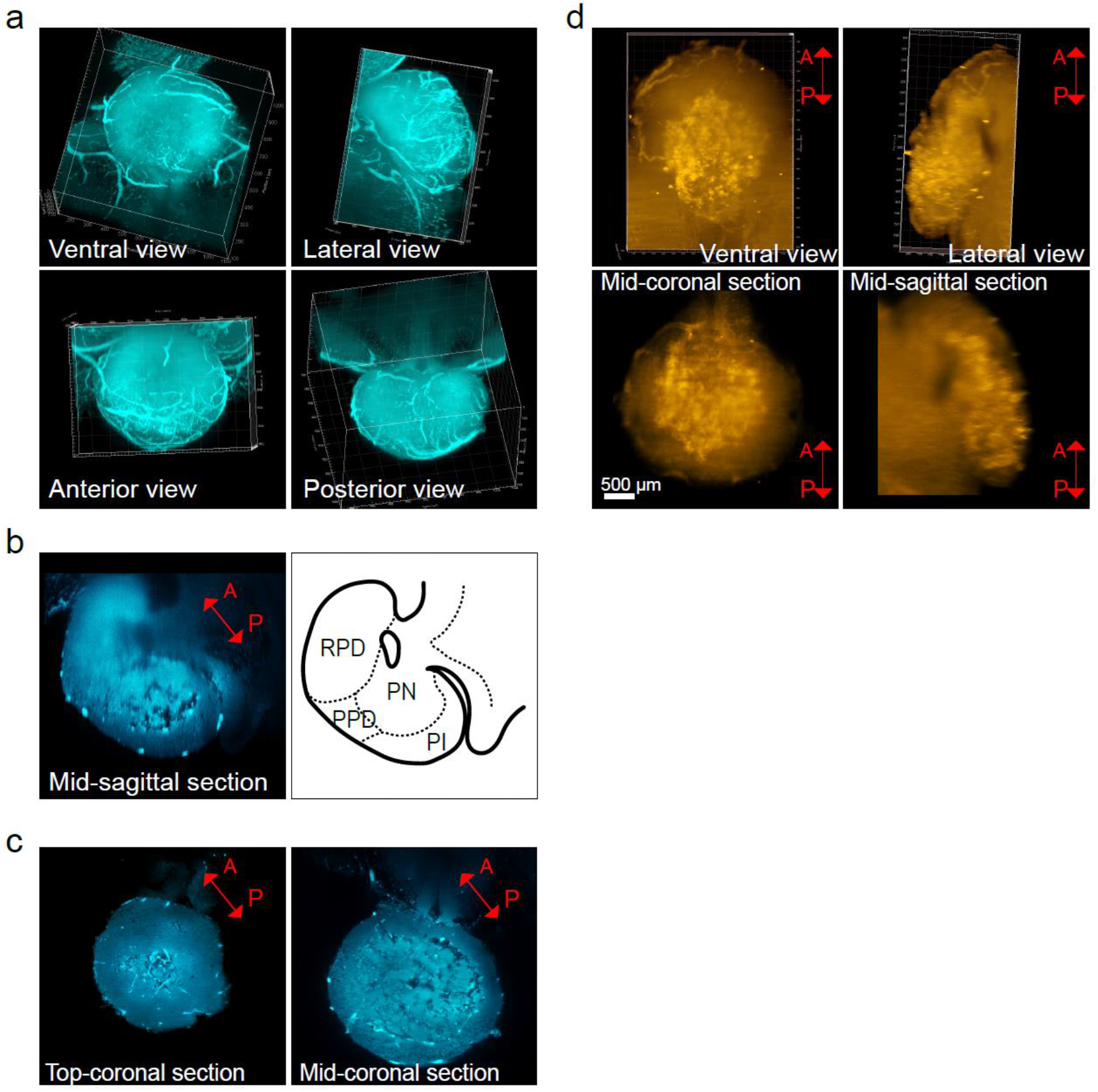
Anatomy of the turquoise killifish pituitary gland. **a** Three-dimensional features of the killifish hypophysis. The transparent killifish brain was stained with DAPI, a nuclear dye. **b** A mid-sagittal section and illustration defining the internal structure of the killifish hypophysis. **c** A coronal section of the killifish hypophysis. Rostral pars distalis (RPD), Proximal pars distalis (PPD), and Pars intermedia (PI), Pars intermedia (PI) and Pars nervosa (PN). **d** Mid-sagittal and coronal sections of the VIP-stained hypophysis. Red arrows indicate the position of the brain; the capital letters “A and B” are abbreviations for anterior and posterior, respectively.

### Anatomy of the turquoise killifish pineal gland

The other important neuroendocrine organ in the brain is the pineal gland. The killifish pineal gland protruded out from the brain in a spherical shape, and then extended a tail into the middle of the telencephalon (Figure 4a, Supplemental movie 3). The coronal section of the pineal gland contained dozens of elliptically-shaped cell arrays (Figure 4a). These globular cell arrays were covered by another layer of cells that were involuted inwards to surround cell arrays, starting from about the middle of the pineal gland and forming an inverted and tailed pocket shape. This outer cell layer section of the pineal gland was located nearest to the telencephalon and the head of the pineal gland placed with optic tectum and habenula (Figure 4a).

**Fig. 4.**
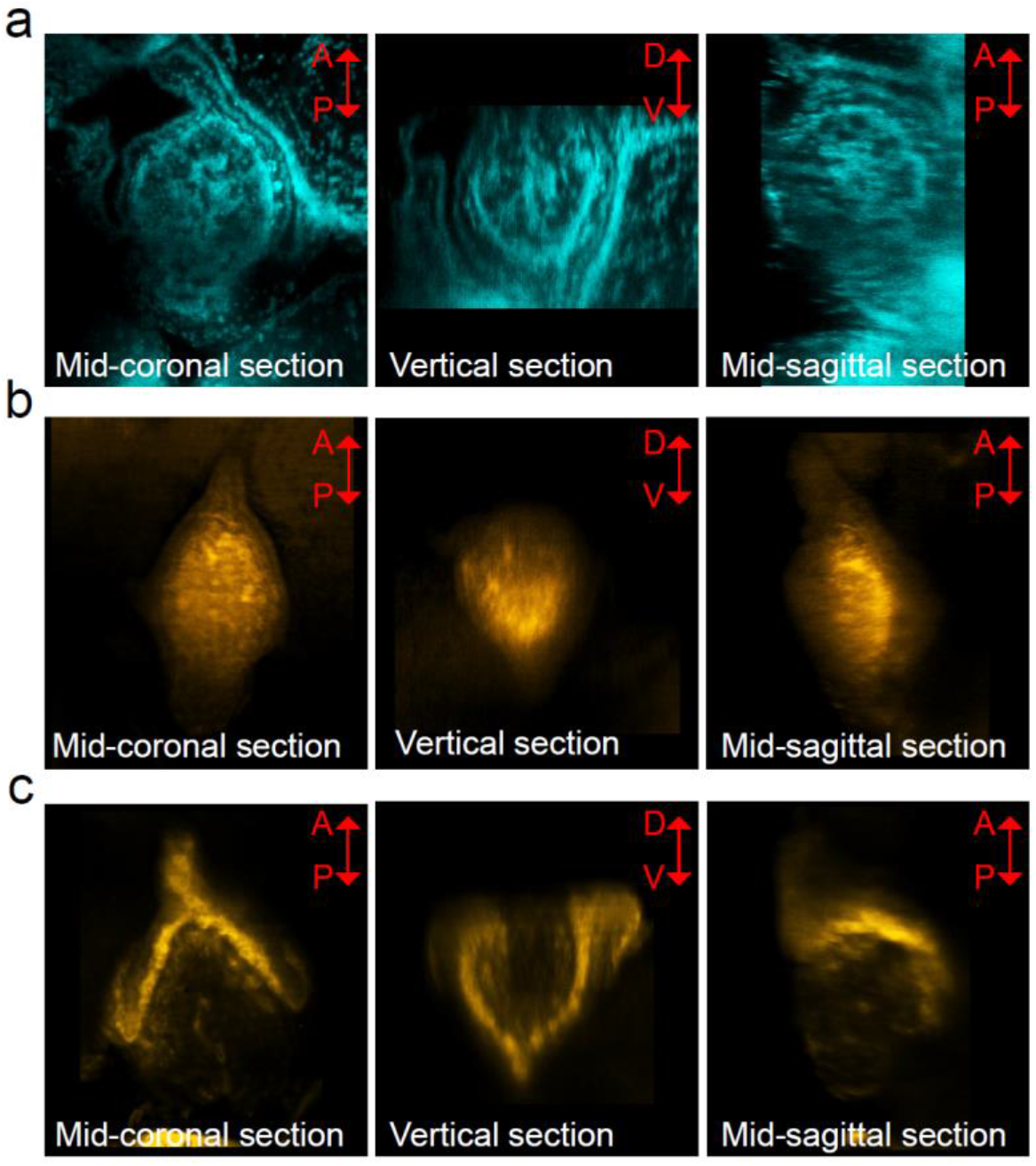
Anatomy of the turquoise killifish epiphysis. **a** Sections of the killifish epiphysis stained with DAPI. **b** Sections of the killifish epiphysis stained with Vimentin. **c** Sections of the killifish epiphysis stained with GFAP. Red arrows indicate position of the brain; the capital letters A, P, D, and V are abbreviations for anterior, posterior, dorsal, and ventral, respectively.

The mammalian pineal gland consists mainly of pinealocytes, as well as astrocytes and microglia (Ibanez Rodriguez et al., 2016; Jiang-Shieh et al., 2003; Moller and Baeres, 2002). To identify microglia and astrocytes, we used the characteristic markers vimentin and glial fibrillary acidic protein (GFAP), respectively (Graeber et al., 1988; Wohl et al., 2011; Zhang et al., 2019). Both vimentin and GFAP were expressed in the killifish pineal gland (Figure 4b, 4c). Vimentin was detected specifically in the proximal region of the pineal gland (Figure 4b, Supplemental movie 4), whereas GFAP localized preferentially to the outer cell layer of the pineal gland, covering half of the globular cell arrays (Figure 4c, Supplemental movie 5).

## Discussion

The neuroendocrine system plays a critical role in harmonizing physiology with behavior in response to external stimuli. This system governs a wide range of biological processes including growth, reproduction, and metabolism. It is well known that these biological processes dramatically decline with aging. Due to the ethical issues surrounding the use of humans for experimental studies, as well as the long lifespan of mammalian model organisms, a new vertebrate model of aging is warranted to satisfy these requirements. The turquoise killifish is a novel model organism for studying the aging process because of its short life span and highly conserved aging phenotypes, including both visual and molecular alterations (Kim et al., 2016). In this study, the neuroendocrine system was described in the turquoise killifish brain. The pituitary gland is the master organ of the neuroendocrine system in killifish pituitary, which is located in the ventral brain between the optic nerves and hypothalamus. Another important neuroendocrine organ, the pineal gland, is located in the dorsal brain between the telencephalon and optic tectum. Their localizations are consistent with other teleost fish species (Trudeau and Somoza, 2020). Taking advantage of the short lifespan and conserved neuroendocrine system, the turquoise killifish is a good model for elucidating an age-dependent dysregulation of the vertebrate neuroendocrine system.

In this study, the neuropeptide VIP was specifically and strongly expressed in the PN of the killifish pituitary gland. VIP is known to be secreted by the RPD (Lam, 1991). After synthesis by the hypothalamus, VIP is hypothesized to travel via the blood stream through the RPD, and is then stored in the PN. VIP is also known to play crucial roles in regulating the circadian clock (Colwell et al., 2003; Hamnett et al., 2019; Vosko et al., 2015; Vosko et al., 2007) because it is strongly localized in the suprachiasmatic nucleus of the mammalian brain. Interestingly, VIP had accumulated in the pituitary, rather than the SCN, of the turquoise killifish brain. There is mounting evidence that the circadian clock system also governs diverse physiologies of the turquoise killifish (Lucas-Sanchez et al., 2011; Lucas-Sanchez et al., 2013; Lucas-Sanchez et al., 2015). Furthermore, the pineal gland is the main organ that secretes melatonin, which is critical for the rhythmic behavior of fish, such as sleep–wake cycles, and other circadian-clock-related activities. Taken together, this study elucidated the key structures of the neuroendocrine system in the turquoise killifish brain, which may be a key site ofr circadian regulatory network.

## Materials and methods

### Fish husbandry and sampling

The short-lived turquoise killifish strain (GRZ-AD) was used and maintained as previously described (Dodzian et al., 2018). The fish room was kept in a 12/12 h light/dark cycle. Fish were fed twice a day at 1 and 8 h after lights on every day.

Every fish in this study was euthanized by supplying 1.5 g/L of ethyl 3-aminobenzoate methanesulfonate (MS-222, E10521, Sigma-Aldrich) to the fish tank water. When fish gill movement ceased, dissections were performed.

The animal husbandry and experiments in this study were carried out in accordance with the animal care and use protocol that is reviewed and approved by the Institutional Animal Care and Use Committee at Daegu Gyeongbuk Institute of Science and Technology, Republic of Korea (approval number: DGIST-IACUC-17103001-00).

### Gross anatomy of the turquoise killifish brain

The heads of young adult fish (8–10 weeks old) were fixed in 4% paraformaldehyde (PFA) overnight. Fish heads were carefully dissected under a stereomicroscope. Brain images were acquired in a 4% PFA solution using a stereomicroscope (M80, Leica Microsystems).

### Nissl staining of killifish brain sections

Whole fish were fixed in 4% PFA overnight. Dozens of intact brains were carefully dissected under a microscope and further fixed before sectioning. Fixed brains were embedded in 2.5% agarose and sectioned with a Vibratome (VT1200S, Leica) at a thickness of 100 μm. Sections were transferred onto slide glasses and dried for 10 min. Sections were stained for 5 min in Nissl stain solution (0.1% cresyl violet (Sigma C-1971), 0.1% glacial acetic acid) and rinsed in distilled water for 5 min. Subsequently, sections were dehydrated in an ethanol series (50%→70%→95%) and de-stained in 100% ethanol until the sections exhibited the desired amount of staining. Stained sections were mounted with mounting medium (Fisher Chemical^TM^ Permount^TM^ Mounting Medium, SP15-100; Fisher Scientific). Section images were acquired using a stereomicroscope (M80, Leica Microsystems).

### Whole brain immunostaining

Several young (6-week-old, after sexual maturation) and old (14-week-old, around median lifespan) brains were carefully dissected under a microscope and then fixed in 4% PFA overnight. Whole brains were cleared using a tissue immunostaining solution (Binaree Immuno Staining^TM^ Kit for Brain, BINAREE), according to the manufacturer’s protocol. Cleared brains were stained with DAPI (D9542, Sigma-Aldrich) and GFAP-Alexa 647 (ab194325, abcam), Vimentin-Alexa 647 (ab195878, abcam), or VIP-Alexa 647 (bs-0077R-A647, Bioss) antibodies. The killifish GFAP, Vimentin, and VIP proteins showed 69%, 68%, and 46% of amino acid sequence identity, respectively, with those of human. Cleared and stained brains were embedded in 2% low-melting-point agarose in capillaries with an inner diameter of 2 mm. The embedded brains were tiled into 9–30 regions to cover the whole brain and imaged using a 20× objective lens (W Plan-APOCHROMAT 20, Zeiss) on a Light Sheet microscope (Lightsheet Z.1, Zeiss). Stacked images were converted (Imaris File Converter x64.9.2, Bitplane) and displayed using Imaris (Imaris x64 7.6.0, Bitplane).

### Glossary for brain sections

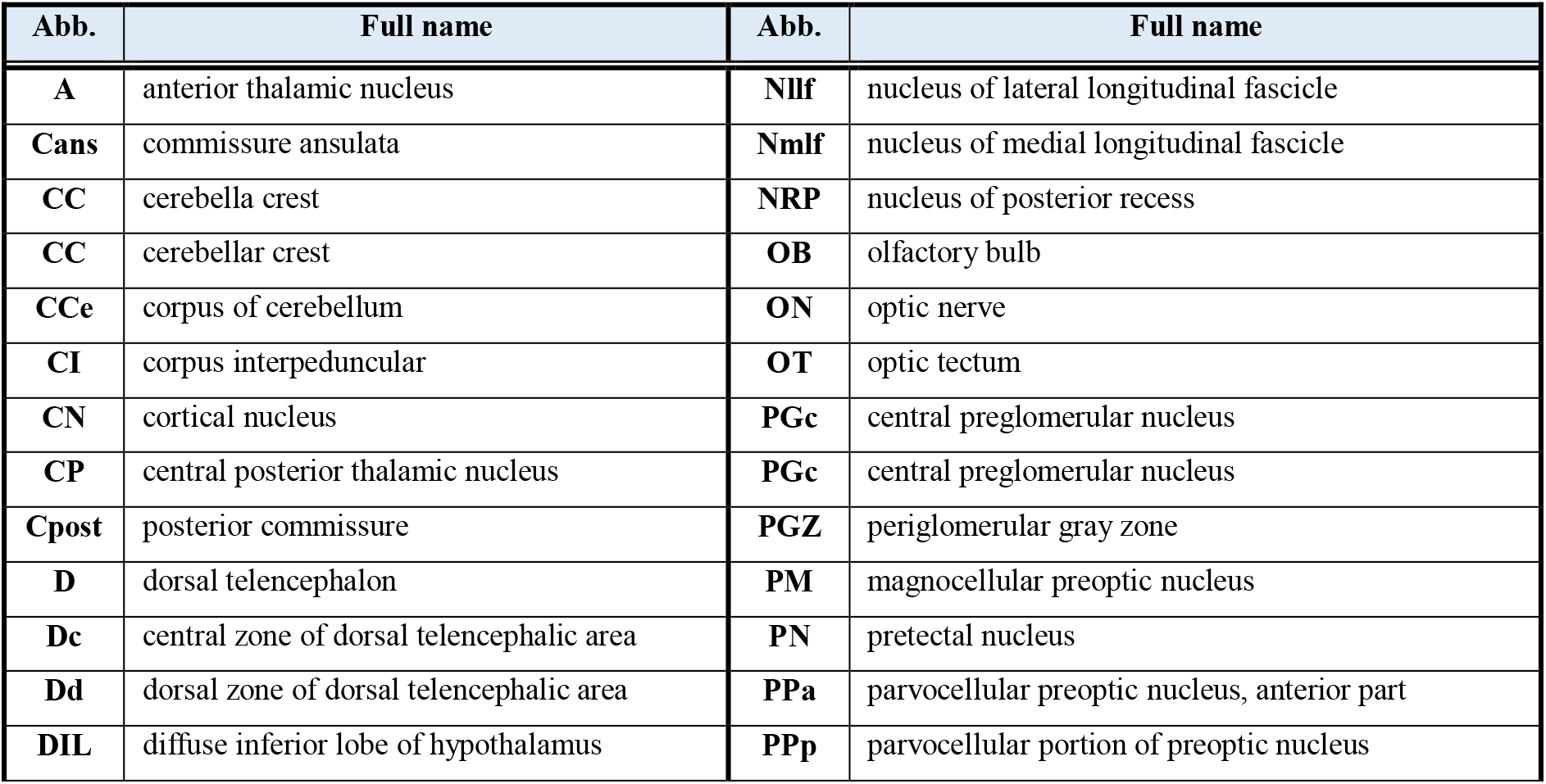
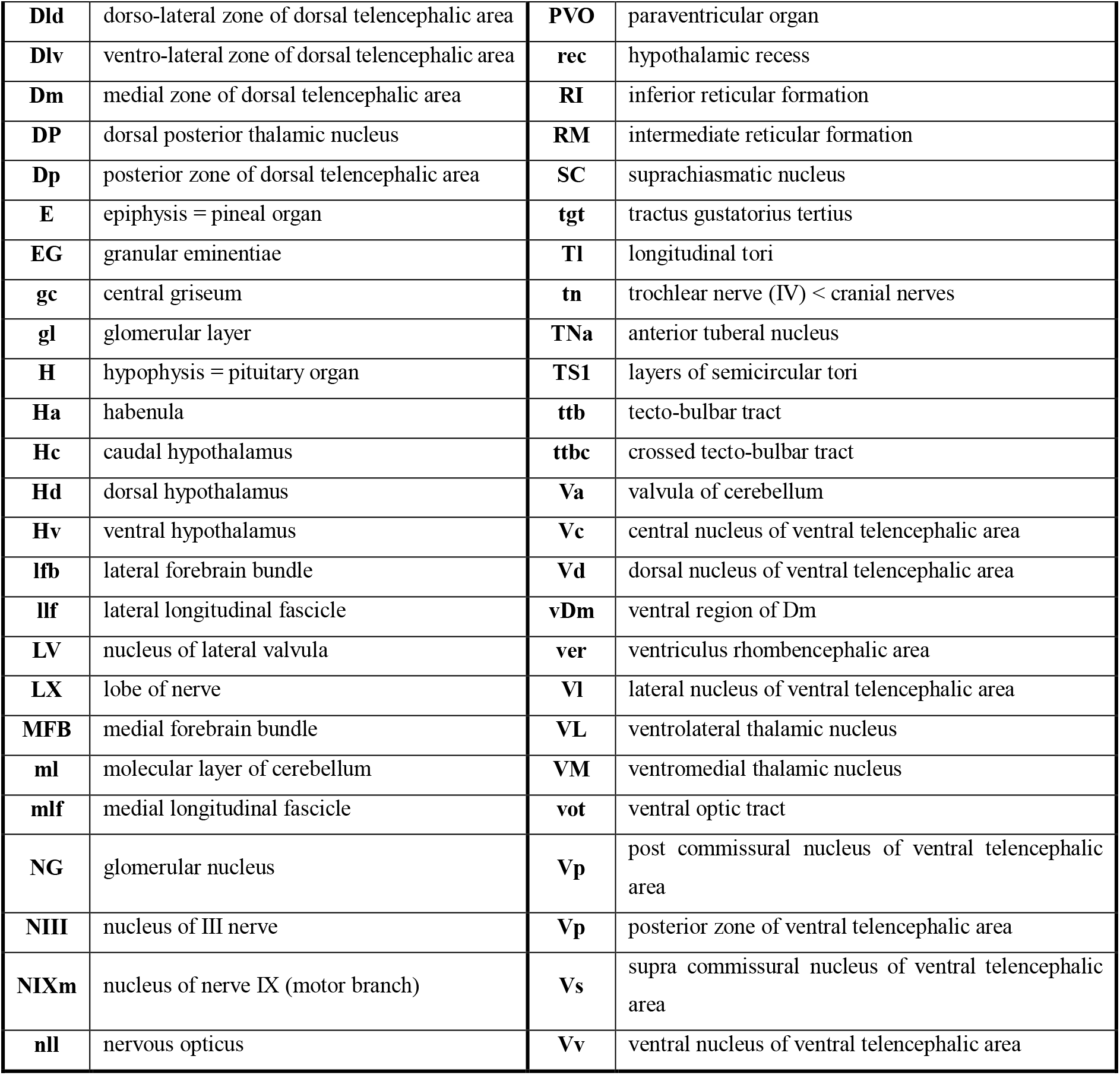

## Acknowledgments

We thank Koichi Kawakami (NIG, Japan) for his helpful suggestion of staining protocols and experimental design discussion.

## Author contributions

YK conceived the project. ED and YK performed the experiments. YK wrote the manuscript. Both authors read and adjusted the manuscript.

## Data and materials availability

All data is available in the main text or the supplementary materials.

## Supplementary Materials

Supplemental Figures 1–3

Supplemental Movies 1–5

