## Supplemental information for "Anatomical identification of the neuroendocrine system in the *Nothobranchius furzeri* brain"

Eunjeong Do[1, †] and Yumi Kim[1,†,*]

This Supplemental information contains Supplemental figures 1 to 3.

Supplemental movies 1 to 5 are provided as separate files.

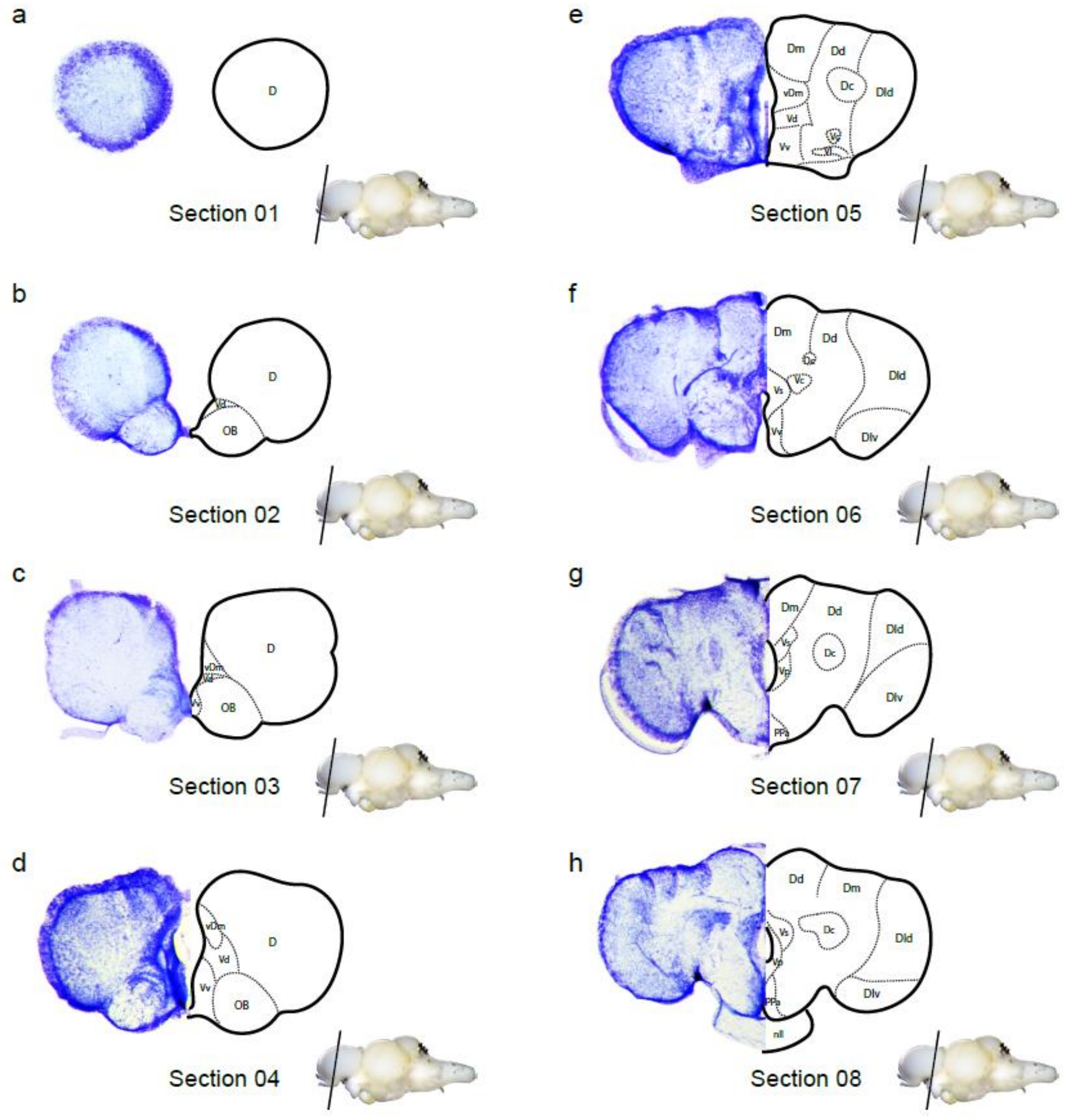


**Supplemental figure 1 Cross-sections of the telencephalic region of the turquoise killifish brain**

**a-e** Sections depict olfactory bulbs of the ventral telencephalic region **f–h**. Sections depict the hypophysis on the ventral brain.

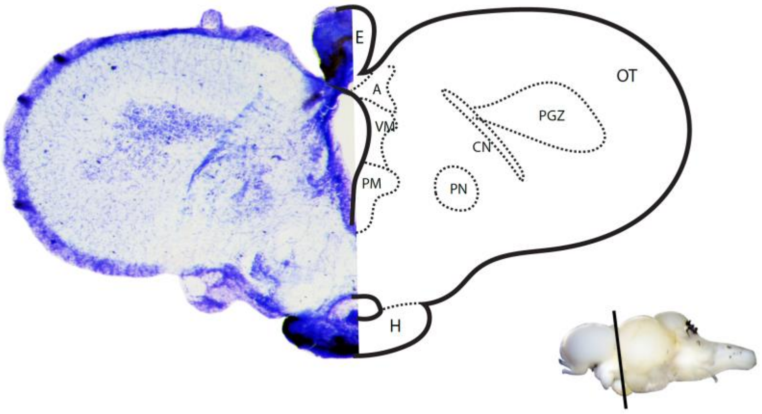


**Supplemental figure 2 The turquoise killifish brain section containing both hypophysis and epiphysis**

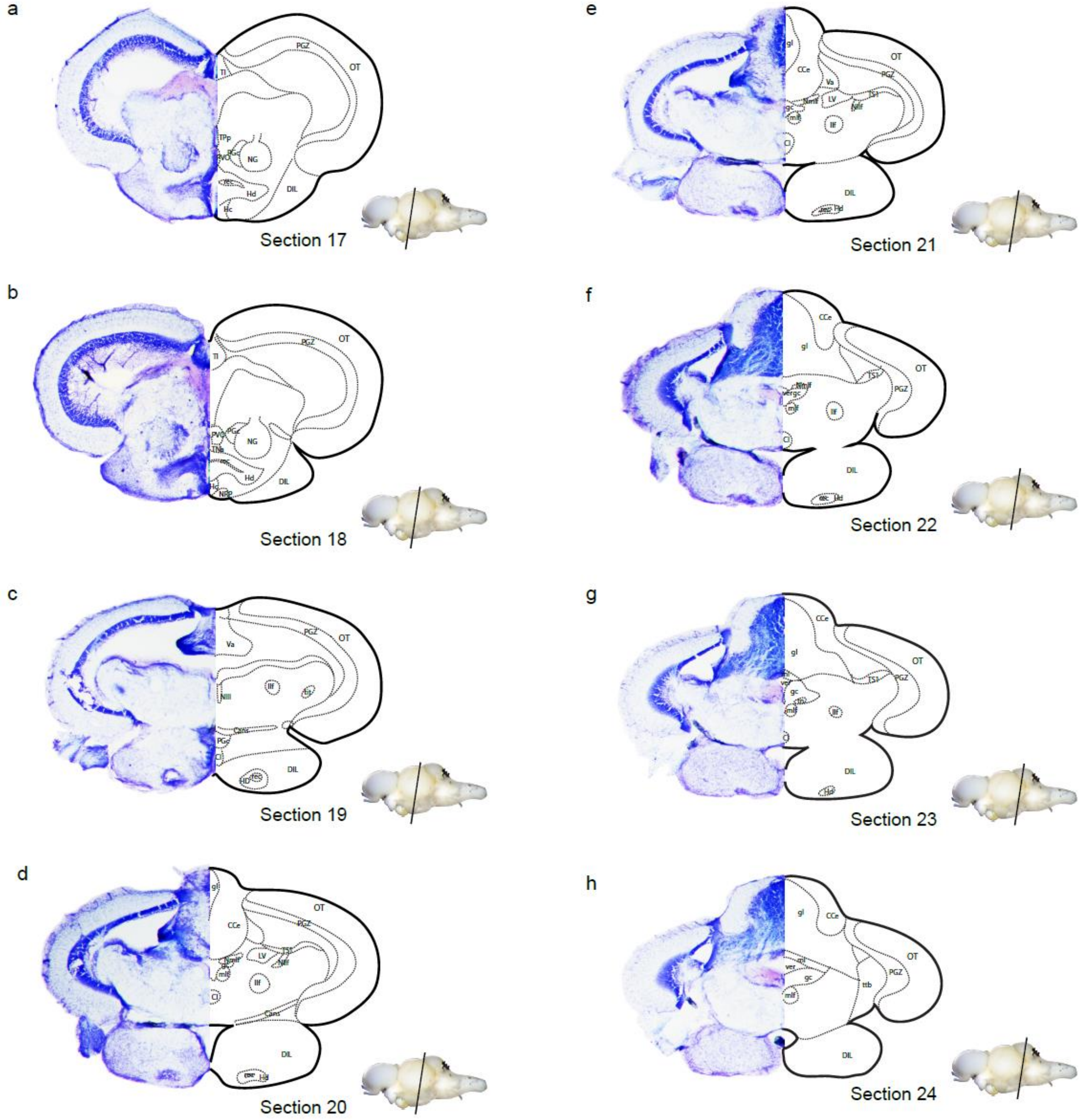
a
Section 17
b
Section 18
c
Section 19
d
Section 20
e
Section 21
f
Section 22
g
Section 23
h
Section 24

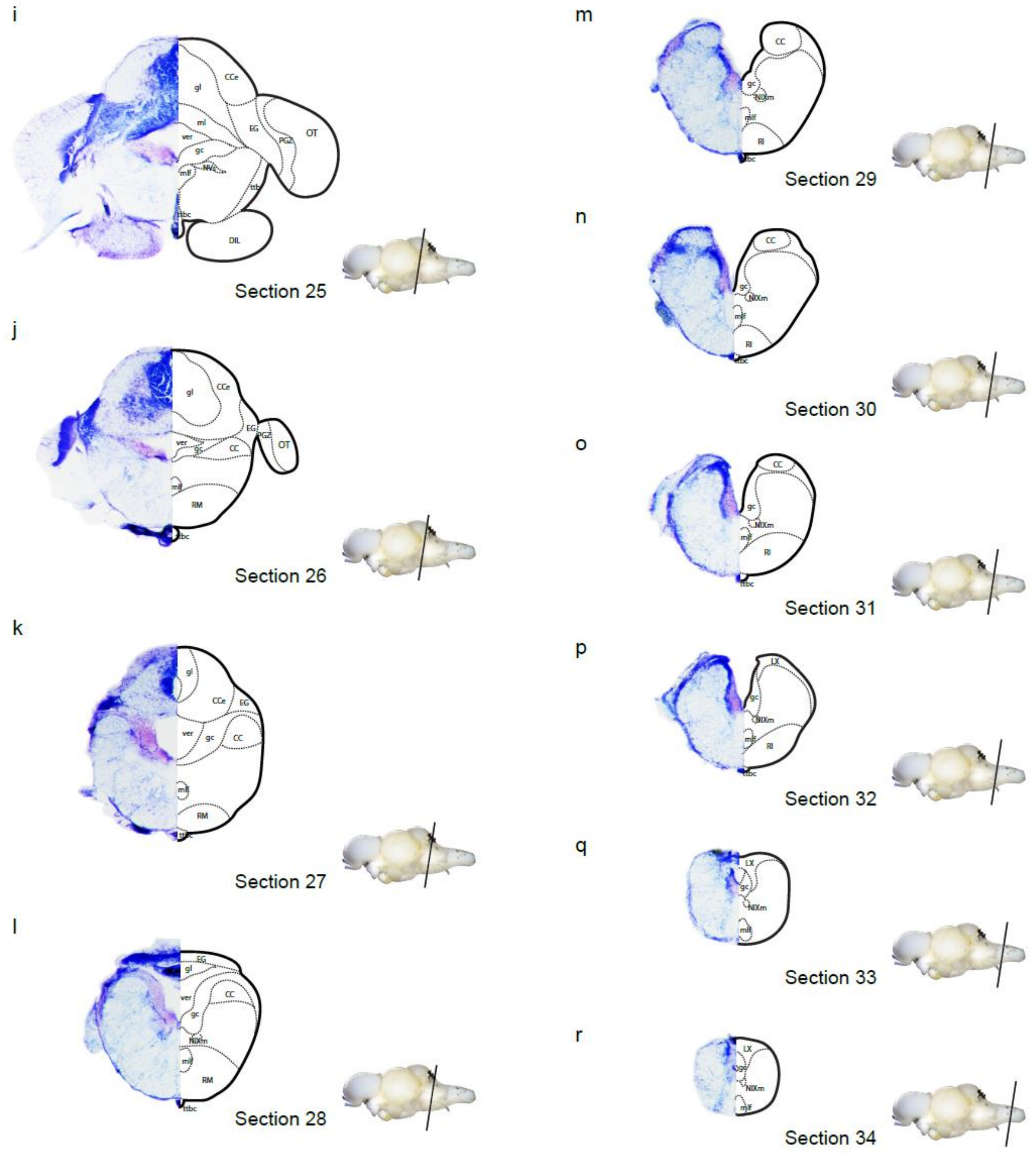


**Supplemental figure 3 Cross-sections of the killifish brain**

**a–e** Sections depict the optic tectum and diencephalic regions. **f–k** Sections contain the cerebellum. **l–r** Sections of the rhombencephalic area

**Supplemental movie 1 Three-dimensional structure of the killifish hypophysis stained with DAPI**

**Supplemental movie 2 Three-dimensional structure of the killifish hypophysis stained with VIP**

**Supplemental movie 3 Movie of the killifish epiphysis optical sections from the dorsal brain and nuclei stained with DAPI**

**Supplemental movie 4 Three-dimensional structure of the killifish epiphysis stained with Vimentin**

**Supplemental movie 5 Three-dimensional structure of the killifish epiphysis stained with GFAP**
